# Estimation of Absolute Protein–DNA Binding Free Energy using Streamlined Geometric Formalism

**DOI:** 10.64898/2026.02.24.707754

**Authors:** Shreya Mukherjee, Diship Srivastava, Niladri Patra

## Abstract

Protein-DNA complexes are involved in vital cellular functions like gene regulation, replication, transcription, packaging, rearrangement, and damage repair. In this work, streamlined geometric formalism for computing the absolute binding free energy was used to obtain chemical accurate *in silico* estimation of binding free energy of three Protein-DNA complexes. Additionally, molecular interactions between Protein and DNA involved hydrogen bonds, electrostatic, van der Waals, and hydrophobic interactions. Using this formalism, researcher can obtain the absolute binding free energy for a Protein-DNA complex with remarkable accuracy and modest computational cost.

---

Deoxyribonucleic acid (DNA) is a biological polymer that contains the fundamental genetic information required for development, function, growth, and reproduction of living organism on earth, as highlighted by its inclusion in the central dogma of molecular biology [1]. Vital cellular functions like gene regulation, replication, transcription, packaging, rearrangement, and damage repair are modulated by binding of DNA to the protein resulting in the formation of Protein-DNA complexes [2–4]. Given the significance of the complex, any aberration like DNA mutation, protein misfolding, and environmental stress, can lead to significance degradation of cell normal functions and in extreme case, cell death. One of the well know example exhibiting the importance of Protein-DNA complex can be seen in the case of p53 tumor suppressor protein p53 [5]. Missense mutations in *TP53* gene induce mutations in DNA binding region of p53, resulting in ineffective binding with canonical p53 target genes, and has been attributed to enhanced tumor progression and neoplastic transformation in cancer cells.

Sequence-specificity protein–DNA binding is required for accessing genetic information during typical cellular function and on the basis of sequence specificity, proteins can be classified into three types, namely specific, multi-specific, and nonspecific [3]. The primary interactions present in the binding regions of a Protein-DNA complex include electrostatic interactions between charged groups, hydrogen bonds between protein and DNA (which may include mediation by bound water molecules), hydrophobic effect, van der Waals forces, and stacking of nucleic bases with each other and with aromatic protein residues. External factors like ionic strength, pH, and counter ion concentration may also affect DNA recognition by the protein.

The ability of two bio-molecules (protein and DNA) to associate and interact with each other is reflected in their absolute binding free energy (ABFE) [6]. Experimental verification of the said binding affinity usually requires a large commitment to the financial and synthesis time. To ease this obstacle substantial work has focused on creating computational techniques for accurate *in silico* prediction of binding free energies [7]. Jayaram et al. exploited the generalized Born equation to obtain the binding free energy of EcoRI Endonuclease-DNA complex and determined that the organization of mobile counterions in the vicinity of A and B form of DNA was responsible for selective preference of DNA form in water and 85% ethanol/water mixture [8]. In the work done Heesch et al., binding free energy of Histone-like Nucleoid Structuring (H-NS) protein with three different nucleotide sequences was determined through Steered molecular dynamics, which resulted in the identification of the high affinity consensus sequence [9]. Recent methods like MELD-DNA,[10], multi site *λ* dynamics[11], and machine learning approaches [12] have provided more accurate values for Protein-DNA binding affinity as compared to the older techniques.

The primary impediments in utilizing the abovementioned methods for computing the binding free energy for Protein-DNA complexes include convergence issues for alchemical methods due to lack of sampling between end states leading to hysteresis, scarcity of benchmark studies, limited data availability for machine learning approaches, and utilization of implicit solvent and “end point” approximation during binding energy calculation [13]. Additionally, previous studies involved small sized DNA fragments, translating these studies to large Protein-DNA complex may not be appropriate. The outcome of most of these studies is the relative binding free energy of Protein-DNA complexes, which are fine for comparing the binding of several DNA sequences but do not provide ABFE. A method which allows for the computation of ABFE for a random sized Protein-DNA complex within chemical accuracy is highly desirable.

One of the method for computing chemical accurate ABFE is geometric route method, which has been successfully applied to several protein-ligand and protein-protein systems [14–17]. It involves a series of potential of mean force (PMF) calculations where the conformational, positional, and orientational degrees of freedom involving bound states of two biomolecules (P and D) are harmonically restrained, thus narrowing the configuration entropy available to the system and alleviating the sampling limitations [6, 14, 15]. In particular, restraints include root mean square deviation (RMSD) of P, D, and P-D interface with respect to a suitable equilibrated structure, relative orientation of P and D (through Euler angles Θ, Φ, and Ψ), and relative positions of P and D (through polar and azimuthal angles *θ* and *φ*), leading to a total of 14 independent simulations in the case of generalized framework.

The drawback of abovementioned geometric route is that due to the degenerate nature of RMSD, the corresponding PMF estimations are hard to converge but can be mitigated through the inclusion of an ergodic sampling algorithm like Gaussian accelerated molecular dynamics (GaMD) [18, 19].

The ergodic sampling algorithm implicitly samples the conformational space of binding partners during PMF estimations, reducing the total number of required independent simulations to six. This combination of ergodic sampling algorithm with geometric route is referred to as streamlined geometric formalism. During this formalism, after a short equilibration, a series of PMF calculations are performed in order (Table 1). In Step 1, PMF estimation with Euler Theta as collective variable (CV) is performed without any restraint. From Step 2 onwards, PMF estimation along the CV involves restraints on CVs from previous steps, with their harmonic center set to the minima of their corresponding 1D PMF. After completion of all the PMF calculations along the Eular, polar, azimuthal angles, and r, ABFE 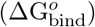 can be computed via the following equations [18]:

**Table 1:** Applied restraints, collective variables, and corresponding free energy contributions during the streamlined geometrical route for computing ABFE. Subscript “o” denotes the contribution of orientational movements. Superscripts “bulk” and “site” denote unbound and bound states.

| Step | Applied Restraint | Collective Variable | Free Energy Contribution |
| --- | --- | --- | --- |
| 1 | - | $\Theta$ | $\Delta G_{\Theta}^{\text{site}}$ |
| 2 | $\Theta$ | $\Phi$ | $\Delta G_{\Phi}^{\text{site}}$ |
| 3 | $\Theta, \Phi$ | $\Psi$ | $\Delta G_{\Psi}^{\text{site}}$ |
| 4 | $\Theta, \Phi, \Psi$ | $\theta$ | $\Delta G_{\theta}^{\text{site}}$ |
| 5 | $\Theta, \Phi, \Psi, \theta$ | $\varphi$ | $\Delta G_{\varphi}^{\text{site}}$ |
| 6 | $\Theta, \Phi, \Psi, \theta, \varphi$ | Distance (r) | $-\frac{1}{\beta} [\ln (S^* I^* C^o)]$ |
| Analytical | - | - | $\Delta G_o^{\text{bulk}}$ |

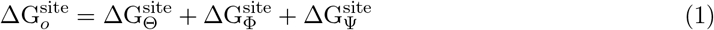

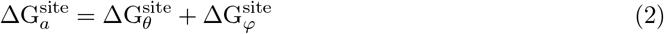

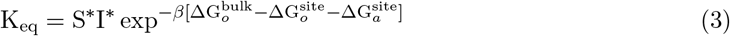

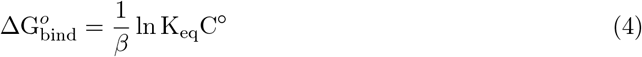

where 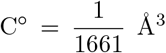 is the standard concentration [15, 18]. Here, 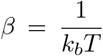, where *k*_*b*_ is the Boltzmann constant and T is the temperature. I^∗^ and S^∗^ denote the separation and surface term respectively. Specific equations for each free energy contribution can found in Supporting Information (SI).

Given the generalized description of streamlined geometric formalism for computing ABFE, it can be hypothesized that same formalism can be used to obtain the ABFE of a Protein-DNA complex. To test this hypothesis, we computed ABFE for following three Protein-DNA complexes using streamlined geometric route:

- **CFP1-CpG Complex**: CXXC domain protein CFP1 bound with non-methylated CpG DNA (PDB: 3QMD) [20]
- **MC1-DNA Complex**: Chromosomal protein MC1 with a 15 base-pair long DNA (PDB: 2NBJ) [21]
- **SopB-DNA Complex**: Centromere-binding protein SopB with an 18 base-pair long DNA (PDB: 3MKW) [22]

During PMF calculation, all the heavy atoms of protein and DNA molecules were used to define the Euler, polar, and azimuthal angles, and distance between the center of mass of the two molecules was used to define CV “r” during separation PMF computation (Figure 1). The individual PMFs were computed using hybrid Gaussian-Accelerated Well-Tempered Metadynamics-Extended Adaptive Biasing Force sampling method (GaWTM-eABF) using Colvar Library and NAMD MD engine [23–25]. In this work, we have defined chemical accuracy such that the difference between computed ABFE and experimental value was less than *k*_*b*_*T*).

**Figure 1.**
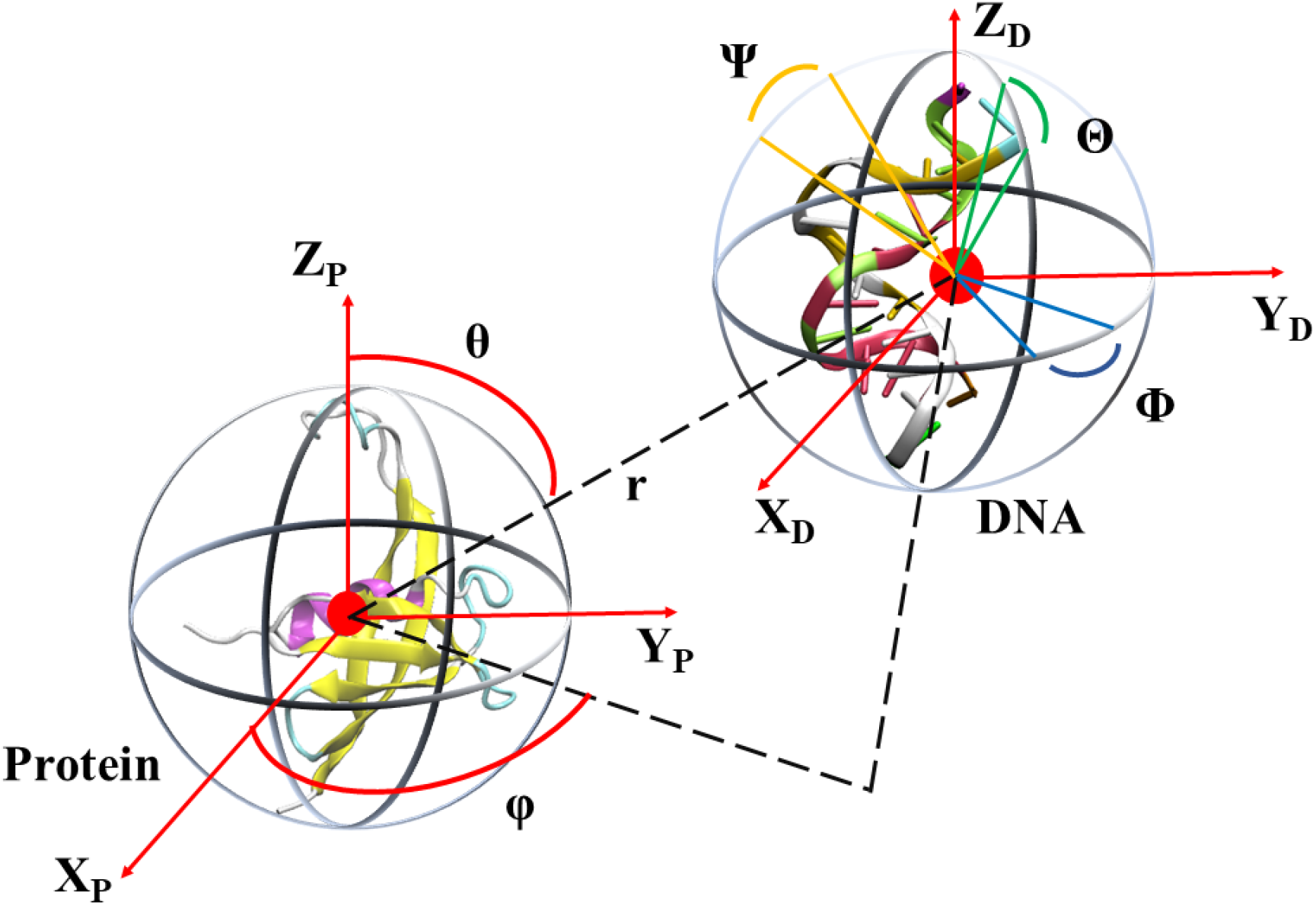
Graphical representation of the Euler, polar, azimuthal angles, and relative separation r between CFP1 (P) and CpG (D) during streamlined geometrical formalism.

In the case of CFP1-CpG Complex, which was smallest of the three Protein-DNA complex under study, we obtained the value of 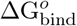 equaling -6.5 *±* 1.4 kcal/mol which matched well with the corresponding experimental value within chemical accuracy (Table 2). Similar results were also observed in the case of MC1-DNA Complex 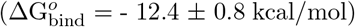 and SopB-DNA Complex 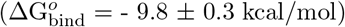, differing only 0.3 kcal/mol and 0.4 kcal/mol from the experimental values. The comparable 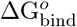 values for the complex’s independent replicas indicated that the formalism was reproducible (Table S2-S4). In all cases, the terms involving 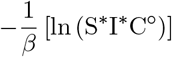 and 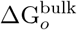 were the primary contributors of 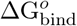. The contribution of orientational and angular terms was marginal but was required for chemical accuracy. Thus, it can be said that chemically accurate the 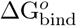 for a Protein-DNA complex can be computed using the streamlined geometric formalism.

**Table 2:** Individual and total free energy terms obtained via the streamlined geometric route for all three systems. Subscript “o” denotes the contribution of orientational movements. Superscripts “bulk” and “site” denote unbound and bound states. All reported ΔG values are mean and standard deviation of three independent replicas and are in kcal/mol. ^∗^ indicates that error metric is less than 0.1 kcal/mol.

|  | CFP1-CpG Complex | MC1-DNA Complex | SopB-DNA Complex |
| --- | --- | --- | --- |
| $\Delta G_{\Theta}^{\text{site}}$ | -0.3* | -0.2* | $-0.3 \pm 0.1$ |
| $\Delta G_{\Phi}^{\text{site}}$ | $-0.4 \pm 0.1$ | $-0.3 \pm 0.1$ | -0.3* |
| $\Delta G_{\Psi}^{\text{site}}$ | $-0.2 \pm 0.1$ | -0.3* | $-0.3 \pm 0.1$ |
| $\Delta G_{\theta}^{\text{site}}$ | -0.2* | -0.2* | -0.1* |
| $\Delta G_{\varphi}^{\text{site}}$ | $-0.2 \pm 0.1$ | $-0.2 \pm 0.1$ | -0.2* |
| $-\frac{1}{\beta} [\ln (S^*I^*C^{\circ})]$ | $-12.0 \pm 1.4$ | $-18.1 \pm 0.6$ | $-15.1 \pm 0.3$ |
| $\Delta G_o^{\text{bulk}}$ | 6.8 | 6.8 | 6.8 |
| $\Delta G_{\text{bind}}^o$ | $-6.5 \pm 1.4$ | $-12.4 \pm 0.8$ | $-9.8 \pm 0.3$ |
| $K_d$ | $11 \pm 2 \mu\text{M}$ [20] | $3.1 \pm 0.6 \text{ nM}$ [21] | $223 \pm 44 \text{ nM}$ [22] |
| $\Delta G_{\text{exp}}$ | -7.0 | -12.1 | -9.4 |

To probe the molecular interactions present in the binding region, we analyzed the MD trajectories of the three complexes (5 independent replicas, each 200 ns long), using ProLIF [26] python library. In all three complex under study, the interactions between protein and DNA were categorized into hydrophobic, van der Waals contact (vdW contact), hydrogen bonds, Pi-cation, and electrostatic interactions.

## CFP1-CpG Complex

In previous studies the crescent-shaped CFP1 CXXC domain was found to be wedged inside the major groove of the non-methylated CpG DNA fragments, leading to the enlargement of the major groove than that of the canonical B-form DNA [20]. The positively charged Lysines and Arginines present on the surface of CFP1 interacted with negatively charged DNA molecule through electrostatic, hydrogen bonds, and vdW contacts. For instance, residues Arg171 and Arg204 were in frequent contacts with DT5’, DC6’, DG7’, DC6’, and DC6 through these interactions. Residues like Ala170 and Ile203 interacted with DG10 and DC6, respectively through hydrophobic contacts. The vdW contacts between residues and bases Gln205-DT8’, Met172-DG7’, and Met194-DA4’ were one of the prominent interactions affecting DNA binding affinity [3]. Pication interactions between bases and residues DC6-Lys202 and DC6-Arg204 were observed in several independent replicas. A number of hydrogen bonds between CFP1 and CpG were present in the complex including but not limited to Arg210-DT5’, DC6’-Arg204, DC6-Ile203, Lys206-DC6’, and Gln205-DG7’ (Figure S ADD FIGURE).

## MC1-DNA Complex

Monomeric nucleoid-associated protein MC1 engages in the genome organization of several *Euryarchaea* by virtue of an atypical compaction mechanism [21]. It is also involved in DNA transcription and cell division and primarily binds to DNA’s minor groove. Positively charged residues Arg4, Lys22, Arg25, Lys30, Lys69, Arg71, and Lys81 were found to be electrostatically bound to the negatively charged DNA backbone atoms. Hydrophobic contacts involved the intercalation of residues Pro72 and Trp74 between between bases DC14 and DA15. vdW contacts between Ile89 and sugar units of DA4, DA5 and DT30 was also noted. Several hydrogen bonds between residues Lys54, His56, Lys86, Lys91 and phophate groups present in the minor groove of the DNA were detected. Additional hydrogen bonds between nucleotides DA4, DC7, DG21, DT22, and DG23 and the side chains of Arg4, Lys22, Gln23, Arg25, and Gln26 were also observed. Pi-cation interactions between DA15-Lys69 and DC14-Arg34 were also present.

## SopB-DNA Complex

The segregation of prototypical F plasmid involve centromere-binding protein SopB, the NTPase SopA and the *sopC* centromere and of which SopB is a multidoamin protein containing all helical DNA-binding domains that are flexibly attached to (*β*_3_ − *α*)_2_ dimer domain [22]. Similar to previous two systems, positively charged Lysines and Arginines (Arg163, Arg190, Lys191, Arg195, Arg219, and Lys226) electrostatically interact with the negatively charged atoms of DNA molecules. The hydrophobic interaction between SopB and DNA involved the amino acids Ile179, Ile192, Ala218, and Ala270. Hydrogen bonds between residues Ser189, Ser217, Ser220, Arg163, Ser220, Arg219, ILe179, Asn178, Ser180, and Thr194 and DNA phophate groups were observed. Additional hydrogen bonds included the donor acceptor pairs Arg190-DG3, Arg190-DG4, Lys191-DG5, Arg194, DG7’, Arg219-DG8’, Ser189-DT6’, Ile192-DT6’, and Lys191-DT6’.

In this work, streamlined geometric formalism was used to obtain the ABFE of three Protein-DNA complexes namely, CFP1-CpG Complex, MC1-DNA Complex, and SopB-DNA Complex.The ABFE values for these complexes was found to be -6.5 kcal/mol, -12.4 kcal/mol, and -9.4 kcal/mol, respectively and differed less than 0.6kcal/mol from the respective experimental values. We also determined that in each system, the primary interactions between protein and DNA involved electrostatic, hydrogen bonds, hydrophobic, vdW contacts and Pi-cation interactions.

For a protein with multiple DNA binding sites, streamlined geometric formalism can be used to accurately provide ABFE and binding constant (using Equation 4) for each binding site and target DNA fragment, which can be compared to the one obtained experimentally and allow the researcher to know the binding site location of target DNA fragment. The protocol described here can be improvement through the application of telescopic solvation solvation scheme during Step 6 (Table 1) [27]. Given that Ribonucleic acid (RNA) is more flexible than DNA, in future work it would be interesting to evaluate the ABFE of a Protein-RNA complex using the streamlined geometric formalism.

## Conflicts of interest

The authors declare no competing financial interest.

## Data Availability Statement

## Acknowledgement

N.P. acknowledges CRG/2020/002308 for funding. S. M. acknowledges DST for awarding the Inspire fellowship (DST/INSPIRE Fellowship/2022/IF220481). S.M. and D.S. acknowledge the usage of HPC cluster present in Indian Institute of Technology (ISM), Dhanbad.

## Associated Content

**For Table of Contents Only**

## Notes

### Competing Interest Statement

The authors have declared no competing interest.

