## Supplemental Figure 1 for "Estimation of Absolute Protein–DNA Binding Free Energy using Streamlined Geometric Formalism"

#### Supporting Information

Shreya Mukherjee[a], Diship Srivastava[a], and Niladri Patra[a*]

[*]

[a]Department of Chemistry and Chemical Biology, Indian Institute of Technology (ISM) Dhanbad, Dhanbad - 826004, India

#### 1 Theory

##### 1.1 Terms appearing in absolute binding free energy ($\Delta G^o_{\mathbf{bind}}$) (Equation 1-4 in main text)

During stepwise potential of mean force (PMF) computations, harmonic restraints were applied on the collective variables of the previous step (See Table 1 in main manuscript), whose general form is given by [1–3]:

$$u_{(.)} = k(\xi_{(.)} - \xi_{0(.)})^2 \tag{1}$$

where k, $\xi_{(.)}$, and $\xi_{0(.)}$ are the force constant, corresponding collective variable, and reference collective variable value, respectively. The individual terms in equation 1 and 2 in main manuscript is given by:

$$\exp^{\beta \Delta G^{\text{site}}_{\Theta}} = \frac{\displaystyle\int_{\text{site}} d\xi \, e^{-\beta U}}{\displaystyle\int_{\text{site}} d\xi \, e^{-\beta [U + u_{\Theta}]}} \tag{2}$$

$$\exp^{\beta \Delta \mathrm{G}_{\Phi}^{\mathrm{site}}} = \frac{\displaystyle\int_{\mathrm{site}} d\xi \, e^{-\beta[U+u_{\Theta}]}}{\displaystyle\int_{\mathrm{site}} d\xi \, e^{-\beta[U+u_{\Theta}+u_{\Phi}]}} \tag{3}$$

$$\exp^{\beta \Delta \mathrm{G}_{\Psi}^{\mathrm{site}}} = \frac{\displaystyle\int_{\mathrm{site}} d\xi \, e^{-\beta[U+u_{\Theta}+u_{\Phi}]}}{\displaystyle\int_{\mathrm{site}} d\xi \, e^{-\beta[U+u_{\Theta}+u_{\Phi}+u_{\Psi}]}} \tag{4}$$

$$\exp^{\beta \Delta \mathrm{G}_{\theta}^{\mathrm{site}}} = \frac{\displaystyle\int_{\mathrm{site}} d\xi \, e^{-\beta[U+u_{\Theta}+u_{\Phi}+u_{\Psi}]}}{\displaystyle\int_{\mathrm{site}} d\xi \, e^{-\beta[U+u_{\Theta}+u_{\Phi}+u_{\Psi}+u_{\theta}]}} \tag{5}$$

$$\exp^{\beta \Delta \mathrm{G}_{\phi}^{\mathrm{site}}} = \frac{\displaystyle\int_{\mathrm{site}} d\xi \, e^{-\beta[U+u_{\Theta}+u_{\Phi}+u_{\Psi}+u_{\theta}]}}{\displaystyle\int_{\mathrm{site}} d\xi \, e^{-\beta[U+u_{\Theta}+u_{\Phi}+u_{\Psi}+u_{\theta}+u_{\phi}]}} \tag{6}$$

$$S^{*} = r^{*2} \int_{0}^{\pi} d\theta \, \sin\theta \int_{0}^{2\pi} d\varphi \, e^{-\beta(u_{\theta}+u_{\varphi})} \tag{7}$$

$$I^{*} = \int_{\mathrm{site}} dr \, e^{-\beta[w(r)-w(r^{*})]} \tag{8}$$

Here $w(r)$ and $r^{*}$ is the PMF along the collective variable and distance to a point far from the binding site ($r^{*} \approx 30$ Å), respectively. $S^{*}$ and $I^{*}$ represents the physical separation of DNA from the binding along the axis defined by the polar and azimuthal angles. $\beta = \frac{1}{k_b T}$, where $k_b$ is the Boltzmann constant and T is the temperature.

#### 1.2 GaWTM-eABF Sampling

In Gaussian-Accelerated Well-Tempered Metadynamics-Extended Adaptive Biasing Force sampling method (GaWTM-eABF) [4], the potential of the system is first modified by the Gaussian accelerated molecular dynamics (GaMD) boost potential, resulting in the reduction of the free energy barrier for all the degrees of freedom present in the system. Then the enhanced sampling along the selected collective variables was performed using WTM-eABF algorithm.

### 2 Methods

#### 2.1 System Preparation Details

The reference structures for the protein-DNA complexes were acquired from RCSB-PDB database for CFP1-CpG Complex (PDB Id: 3QMD) [5], MC1-DNA Complex (PDB Id: 2NBJ) [6], and SopB-DNA Complex (PDB Id: 3MKW) [7]. The PDB structures thus obtained were subjected to the PDB Reader and Manipulator module of CHARMM-GUI to remove crystallographic water, add missing residues, and cleanup the system [8, 9]. Further, H++ server was used to determine the protonation states of titratable residues in each system at pH 7.0 [10, 11]. Leaprc module was employed to solvate the complexes into an orthorhombic waterbox with padding of 10 Å in all directions. Equivalent amounts of individual $Na^+$ and $Cl^-$ ions were introduced to each system to ensure charge neutrality and a concentration of 150 mM (Table S1).

| | CFP1-CpG Complex | MC1-DNA Complex | SopB-DNA Complex |
|---|---|---|---|
| Number of base pairs | 12 | 15 | 18 |
| Number of protein residues | 79 | 93 | 138 |
| Total number of atoms | 19467 | 26699 | 41908 |
| Water Molecules | 5931 | 8045 | 12956 |
| $Na^+$ | 37 | 49 | 77 |
| $Cl^-$ | 23 | 31 | 47 |

Table S1: System details for each of the three systems.

#### 2.2 Computational Settings During Classical Molecular Dynamics

All equilibrium molecular dynamics (MD) simulation was performed using Amber 19 software [12, 13]. Amber ff14SB and bsc1 force fields were used for defining the topology of each system and corresponding files were prepared using AmberTools18's tLeap module [14–16]. Energy minimization was carried out using Steepest Descent in the initial steps and then by Conjugate Gradient method [17]. A gradual NVT heating was conducted from 5K to 310K while applying various positional restraints. Later, NPT equilibration was performed, where a constant temperature of 310 K and pressure of 1 atm was maintained using Langevin thermostat with a collision frequency of 1 $ps^{-1}$ and Berendsen Barostat, respectively [18]. Periodic electrostatics of the systems were maintained by Particle Mesh Ewald (PME) method and the covalent hydrogen bonds were constrained by enabling SHAKE algorithm [19, 20]. During the 500 ns long production run, the time step was set to 2 fs and trajectory was saved every 40 ps. Five independent replicas were performed for each system. During analysis only last 200 ns trajectory for each replica were taken into account.

##### 2.3 Analysis

The visualization of the system was performed using VMD 1.9.4 software [21]. Data plotting and other analysis were performed using in house python and Tcl scripts. A hydrogen bond was assumed to be formed when the distance between hydrogen bond donor (D) and hydrogen bond acceptor (A) was less than 3.5 Å and $\angle \mathrm{DHA} \in [130°, 180°]$.

##### 2.4 Calculation of Interaction Fingerprint

ProLIF [22] was used to obtain the interaction fingerprint between the protein and DNA molecule in each of the three system, using the default criteria for detecting the type of interaction present in the production trajectories. In the bar plots depicting the interaction fingerprint analysis using ProLIF (Figures S9 - S23), the residue sequence numbers in CFP1-CpG Complex, MC1-DNA complex and SopB-DNA Complex were renumbered to start from 1 instead of the residue sequence numbers provided in the PDB files downloaded from RCSB and can be found in Figures S1 - S3. However in the main manuscript and discussion original residue sequence numbers (provided by RCSB) was used.

#### RCSB Protein:

169 198
SARMCGECEACRRTEDCGHCDFCRDMKKFGGPNKI
221
RQKCRLRQCQLRARESYK

#### ProLIF Protein:

1
SARMCGECEACRRTEDCGHCDFCRDMKKFGGPNKI
36 53
RQKCRLRQCQLRARESYK

#### DNA:

1 12
GCCATCGTTGGC

54 65
GCCATCGTTGGC

Figure S1: The residue sequence numbers in CFP1-CpG Complex used during interaction fingerprint analysis using ProLIF.

#### RCSB Protein:

1 31
SNTRNFVLRDEDGNEHGVFTGKQPRQAALKAANRG

SGTKANPDIIRLRERGTKKVHVFKAWKEIVDAPKNRP

81 93
AWMPEKISKPFVKKERIEKLE

#### DNA:

1 15
AAAAACACACACCCA

94 108
AAAAACACACACCCA

Figure S2: The residue sequence numbers in MC1-DNA Complex used during interaction fingerprint analysis using ProLIF.

RCSB Protein:

165

PTSAYERGQRYASRLQNEFAGNISALADAENISRKIIT

195

RCINTAKLPKSVVALFSHPGELSARSGDALQKAFTDK

232

EELLKQQASNLHEQKKAGVIFEADEVITLLTSVLKTS

271

SAS

ProLIF Protein:

1

PTSAYERGQRYASRLQNEFAGNISALADAENISRKIIT

39

RCINTAKLPKSVVALFSHPGELSARSGDALQKAFTDK

EELLKQQASNLHEQKKAGVIFEADEVITLLTSVLKTS

115

SAS

DNA:

1 18

CTGGGACCATGGTCCCAG

116 113

CTGGGACCATGGTCCCAG

Figure S3: The residue sequence numbers in SopB-DNA Complex used during interaction fingerprint analysis using ProLIF.

##### 2.5 BFEE2 System Preparation And Settings

The input files for streamlined geometrical route were generated using BFEEstimator v2.5.0 [1, 23]. For this purpose, the last frame of the replica 1 of equilibrium MD for each system was chosen as the input structure. Since the GaWTM-eABF pipeline used here does not require enhanced sampling of the protein and DNA molecules' RMSD, only files pertaining to the enhanced sampling of Euler, polar, and azimuthal angles were generated. As for the sampling along distance between the center of mass of heavy atoms of protein and DNA, distance colvar was used [24]. During this step, TIP3P water [25] was added to the initial water box in order to expand it by 30 Å accounting for the separation of protein and DNA molecules. During sampling Euler, polar, and azimuthal angles were harmonically restrained using a force constant equaling 0.1 kcal/(mol deg$^2$) (Equation 1, Table 1 in main text). Additionally, during PMF calculation the translation and orientation of the protein were locked using two separate harmonic restraints. Since individual PMF calculations were performed using Colvar module [24] and NAMD MD engine [26], compatible input files were saved.

##### 2.6 MD protocol during Enhanced Sampling Calculations

NAMD [26] MD engine along with Colvar Module was used to perform the enhanced sampling using GaWTM-eABF method and Amber force field compatible settings. The equations of motion were integrated using a time step of 2 fs and under NPT conditions with constant temperature maintained at 310 K using Langevin dynamics and constant pressure set to 1 atm using Nose-Hoover Langevin piston [27, 28] were used. Smooth PME method was used to obtain the long range periodic electrostatic contribution to the force field [19]. SHAKE [20] and SETTLE [29] were used to constrain the covalent bonds involving non-water and water hydrogen bonds, respectively. For each system three independent replicas were performed.

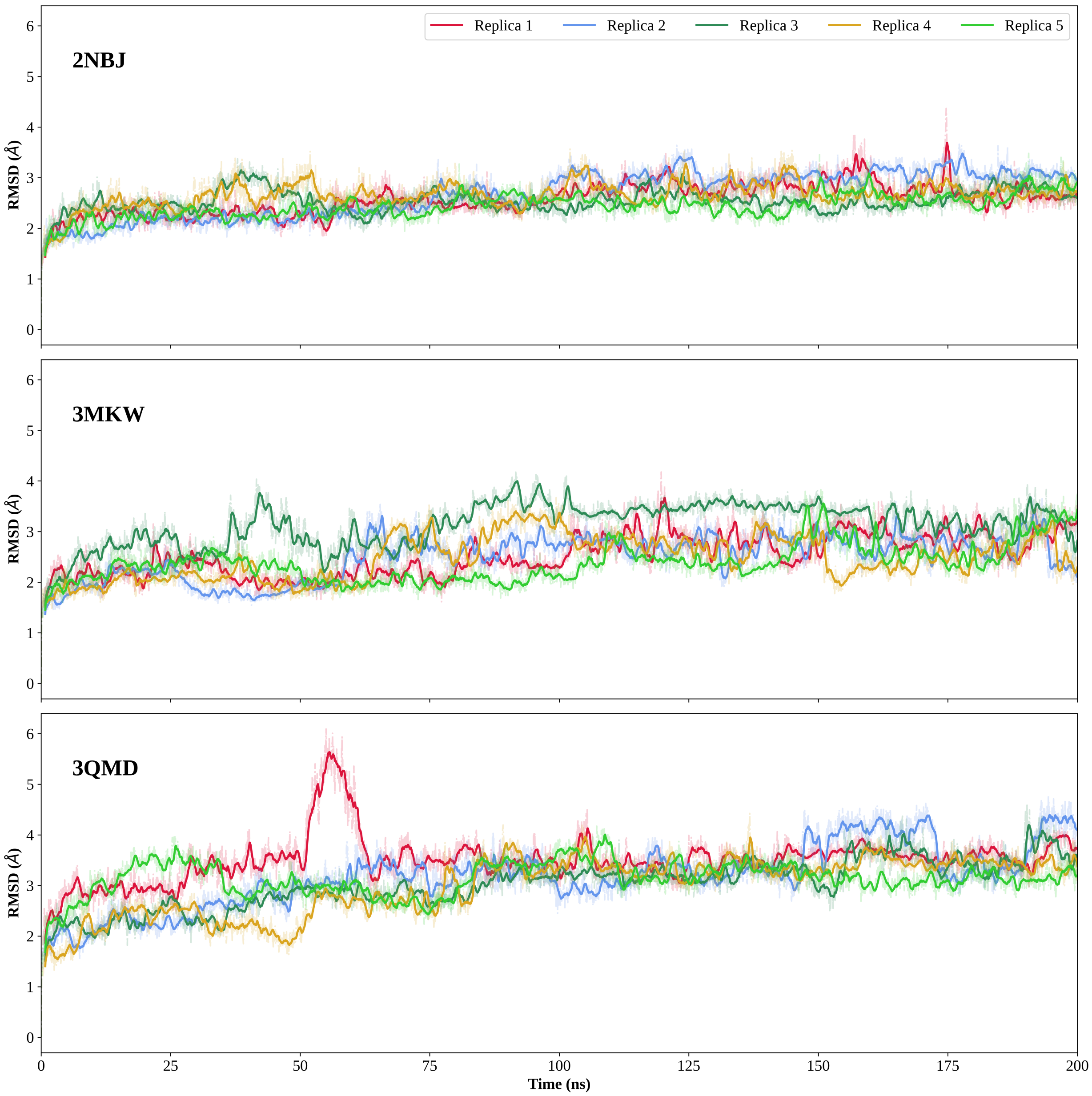


Figure S4: RMSD of protein backbone atoms in the complexes: MC1-DNA Complex (top), SopB-DNA Complex (middle), and CFP1-CpG Complex (bottom), for the last 200 ns of individual replica runs.The darker line plots depict 10 frame running average and shaded areas represent the raw data.

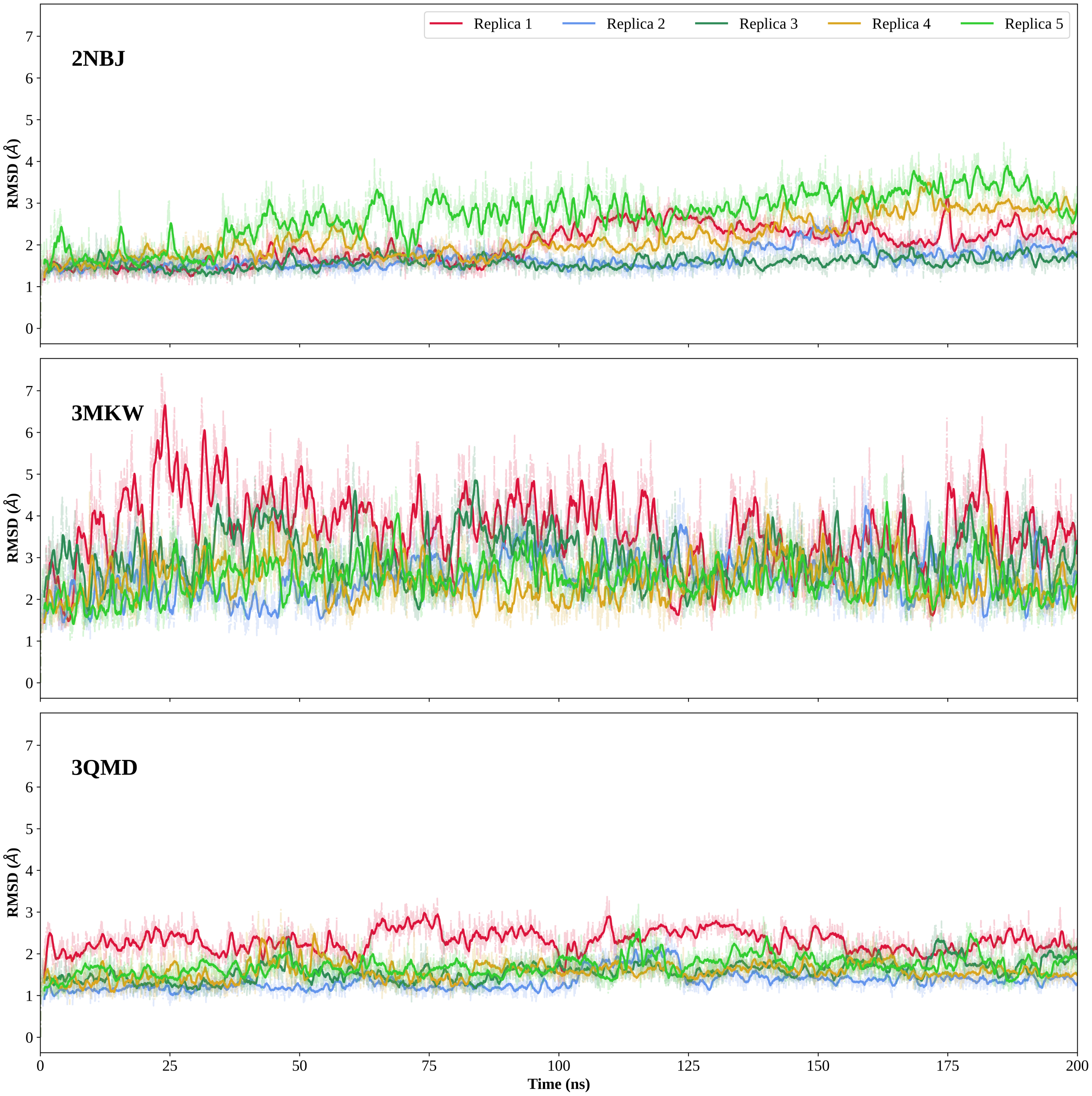


Figure S5: RMSD of DNA backbone atoms in the complexes: MC1-DNA Complex (top), SopB-DNA Complex (middle), and CFP1-CpG Complex (bottom), for the last 200 ns of individual replica runs.The darker line plots depict 10 frame running average and shaded areas represent the raw data.

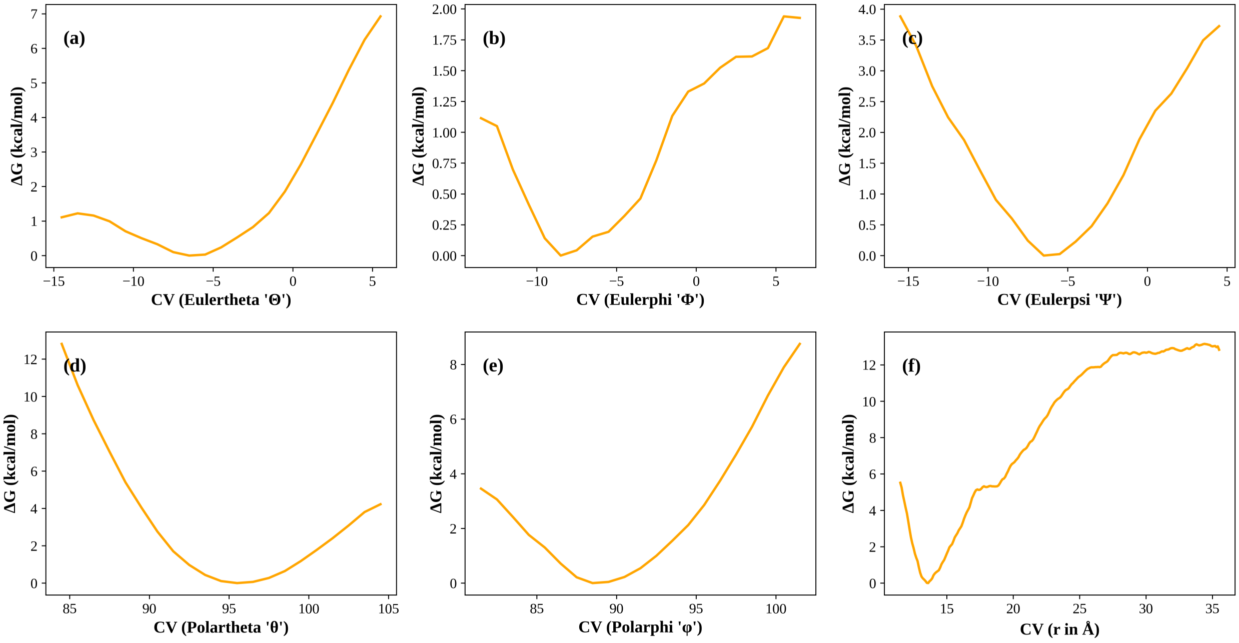


Figure S6: The potential of mean force profiles for different collective variables for the CFP1-CpG Complex.

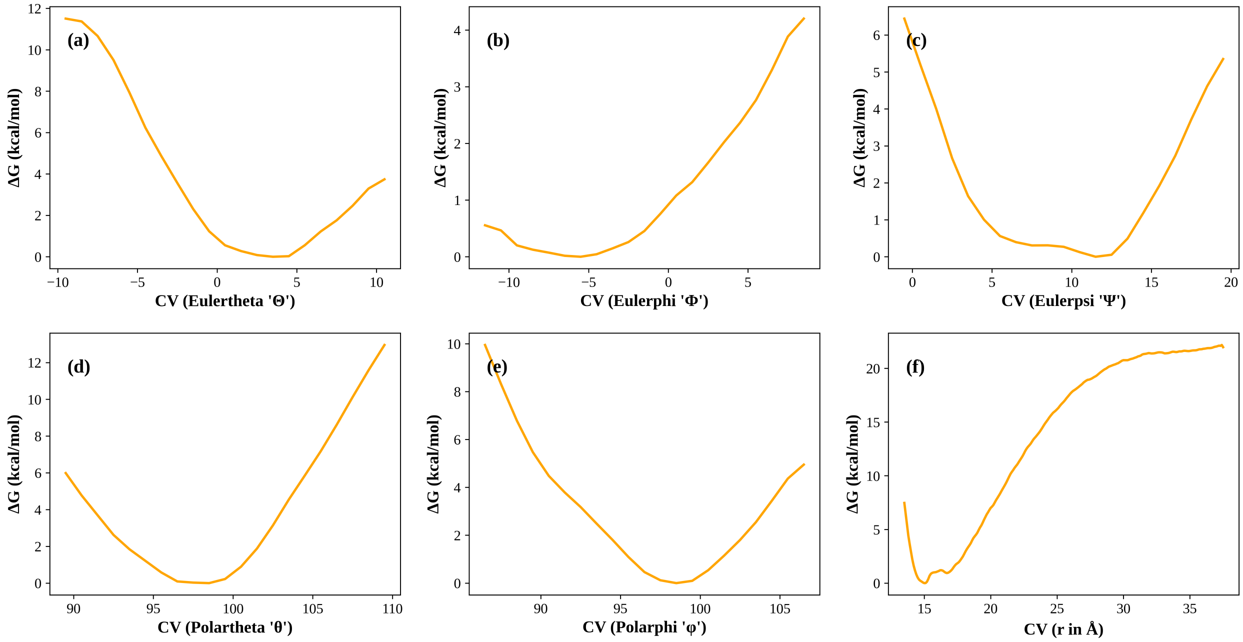


Figure S7: The potential of mean force profiles for different collective variables for the MC1-DNA Complex.

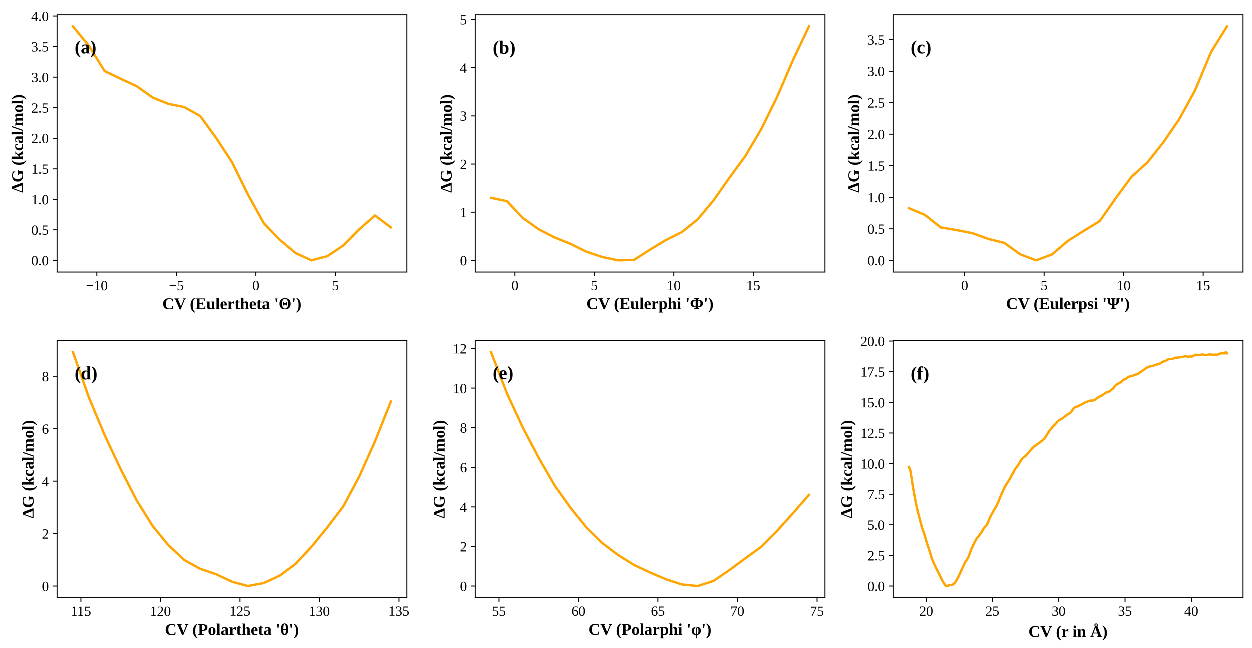


Figure S8: The potential of mean force profiles for different collective variables for the SopB-DNA Complex.

| | Replica 1 | Replica 2 | Replica 3 |
|---|---|---|---|
| $\Delta G_{\Theta}^{\text{site}}$ | -0.26 | -0.30 | -0.30 |
| $\Delta G_{\Phi}^{\text{site}}$ | -0.34 | -0.39 | -0.55 |
| $\Delta G_{\Psi}^{\text{site}}$ | -0.19 | -0.23 | -0.32 |
| $\Delta G_{\theta}^{\text{site}}$ | -0.14 | -0.16 | -0.15 |
| $\Delta G_{\varphi}^{\text{site}}$ | -0.15 | -0.27 | -0.25 |
| $-\frac{1}{\beta}\left[\ln\left(S^{*}I^{*}C^{\circ}\right)\right]$ | -11.76 | -13.13 | -11.15 |
| $\Delta G_{o}^{\text{bulk}}$ | 6.8 | 6.8 | 6.8 |
| $\Delta G_{\text{total}}$ | -6.04 | -7.68 | -5.91 |

Table S2: Individual and total free energy terms obtained via the streamlined geometric route for CFP1-CpG Complex. Subscript "o" denotes the contribution of orientational movements. Superscripts "bulk" and "site" denote unbound and bound states. All reported ΔG values in kcal/mol.

| | Replica 1 | Replica 2 | Replica 3 |
|---|---|---|---|
| $\Delta G_{\Theta}^{\text{site}}$ | -0.21 | -0.13 | -0.17 |
| $\Delta G_{\Phi}^{\text{site}}$ | -0.43 | -0.34 | -0.20 |
| $\Delta G_{\Psi}^{\text{site}}$ | -0.31 | -0.29 | -0.31 |
| $\Delta G_{\theta}^{\text{site}}$ | -0.22 | -0.13 | -0.15 |
| $\Delta G_{\varphi}^{\text{site}}$ | -0.23 | -0.11 | -0.16 |
| $-\frac{1}{\beta}\left[\ln\left(S^{*}I^{*}C^{\circ}\right)\right]$ | -18.37 | -18.54 | -17.34 |
| $\Delta G_{o}^{\text{bulk}}$ | 6.8 | 6.8 | 6.8 |
| $\Delta G_{\text{total}}$ | -12.98 | -12.74 | -11.55 |

Table S3: Individual and total free energy terms obtained via the streamlined geometric route for MC1-DNA complex. Subscript "o" denotes the contribution of orientational movements. Superscripts "bulk" and "site" denote unbound and bound states. All reported ΔG values in kcal/mol.

| | Replica 1 | Replica 2 | Replica 3 |
|---|---|---|---|
| $\Delta G_{\Theta}^{\text{site}}$ | -0.33 | -0.36 | -0.23 |
| $\Delta G_{\Phi}^{\text{site}}$ | -0.27 | -0.27 | -0.30 |
| $\Delta G_{\Psi}^{\text{site}}$ | -0.22 | -0.46 | -0.36 |
| $\Delta G_{\theta}^{\text{site}}$ | -0.13 | -0.15 | -0.16 |
| $\Delta G_{\varphi}^{\text{site}}$ | -0.19 | -0.15 | -0.14 |
| $-\frac{1}{\beta}\left[\ln\left(S^{*}I^{*}C^{\circ}\right)\right]$ | -14.92 | -15.04 | -15.42 |
| $\Delta G_{o}^{\text{bulk}}$ | 6.8 | 6.8 | 6.8 |
| $\Delta G_{\text{total}}$ | -9.26 | -9.63 | -9.82 |

Table S4: Individual and total free energy terms obtained via the streamlined geometric route for SopB-DNA Complex. Subscript "o" denotes the contribution of orientational movements. Superscripts "bulk" and "site" denote unbound and bound states. All reported ΔG values in kcal/mol.

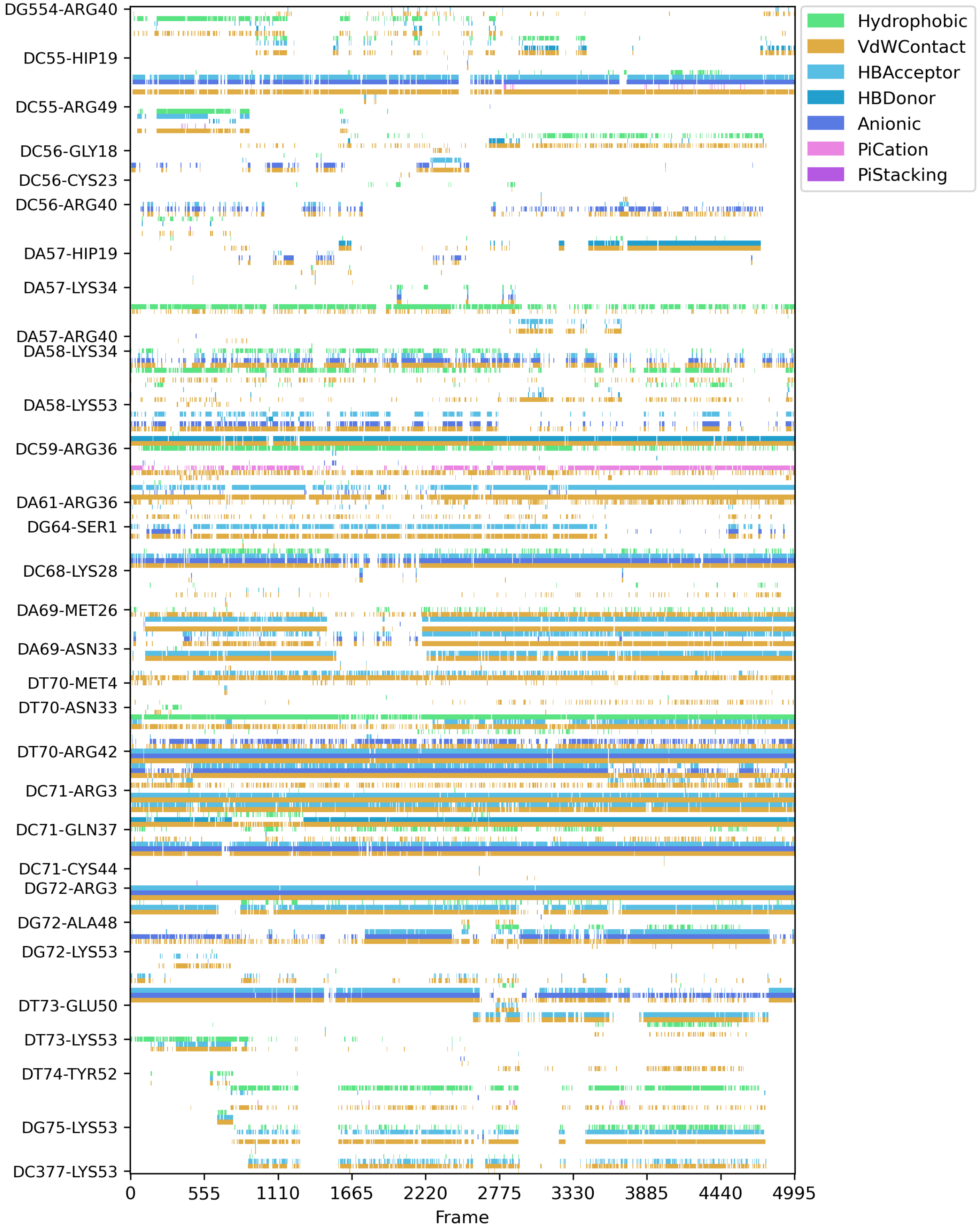


Figure S9: Bar plot depicting the interaction fingerprint between the protein and DNA in CFP1-CpG Complex (Replica 1).

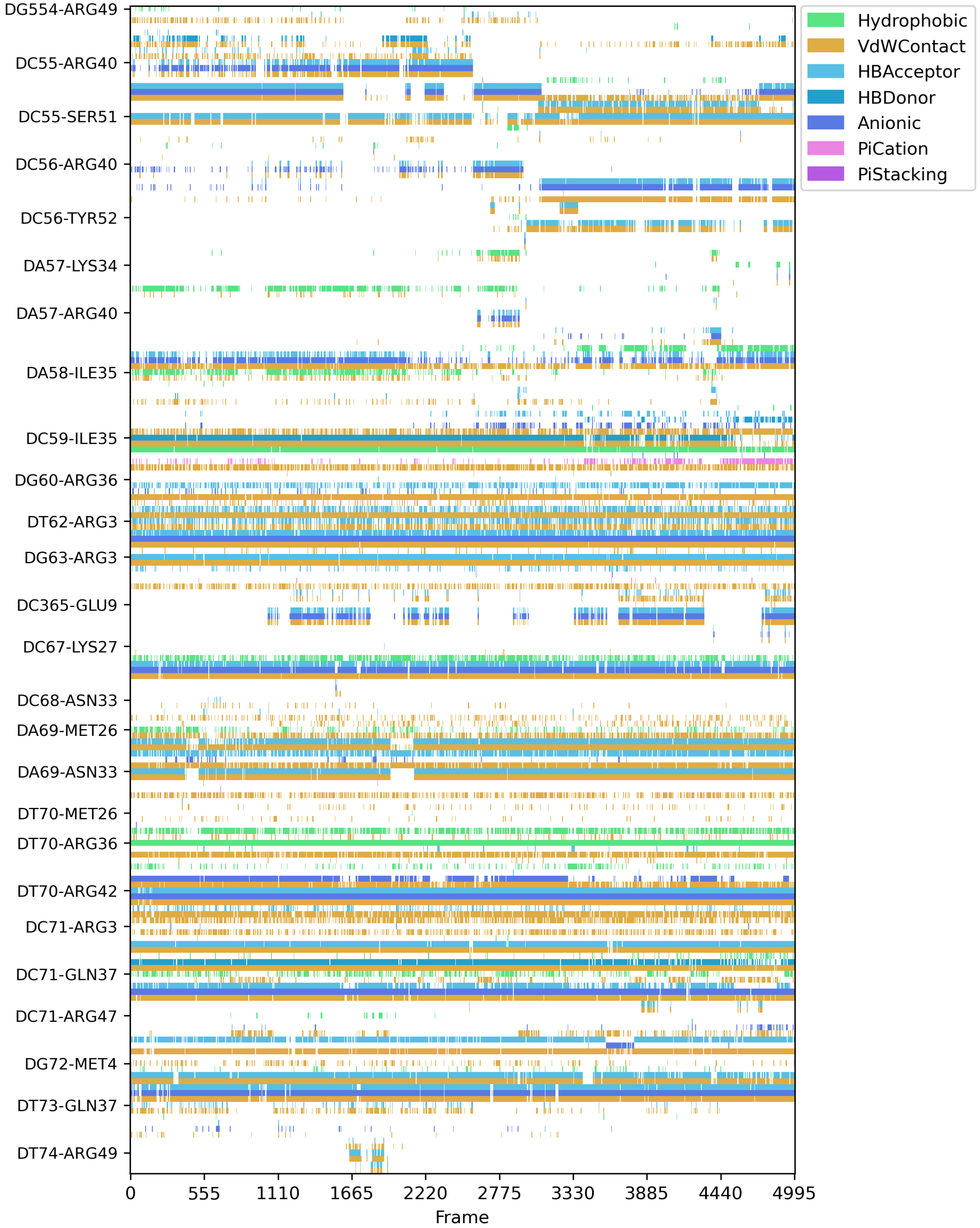


Figure S10: Bar plot depicting the interaction fingerprint between the protein and DNA in CFP1-CpG Complex (Replica 2).

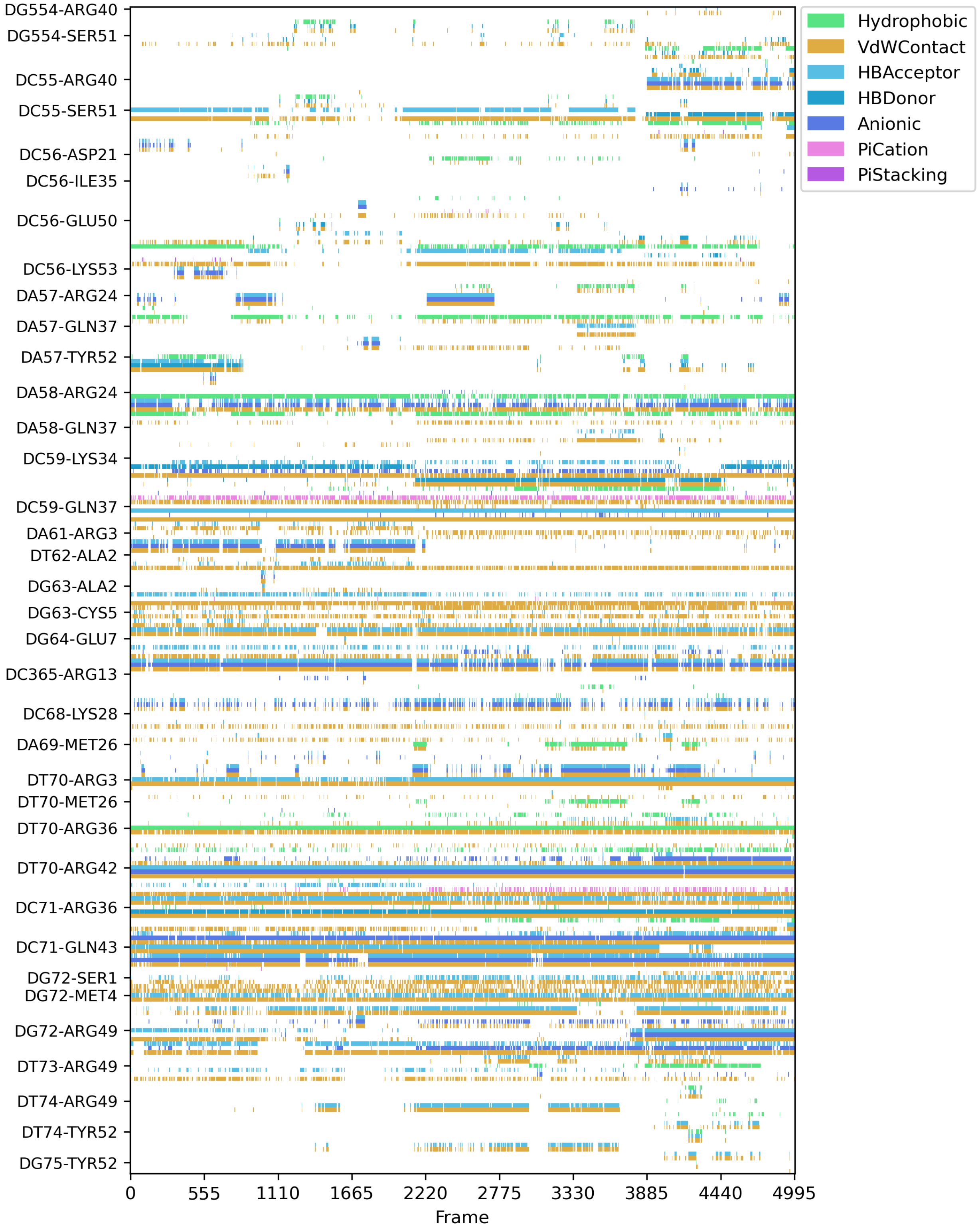


Figure S11: Bar plot depicting the interaction fingerprint between the protein and DNA in CFP1-CpG Complex (Replica 3).

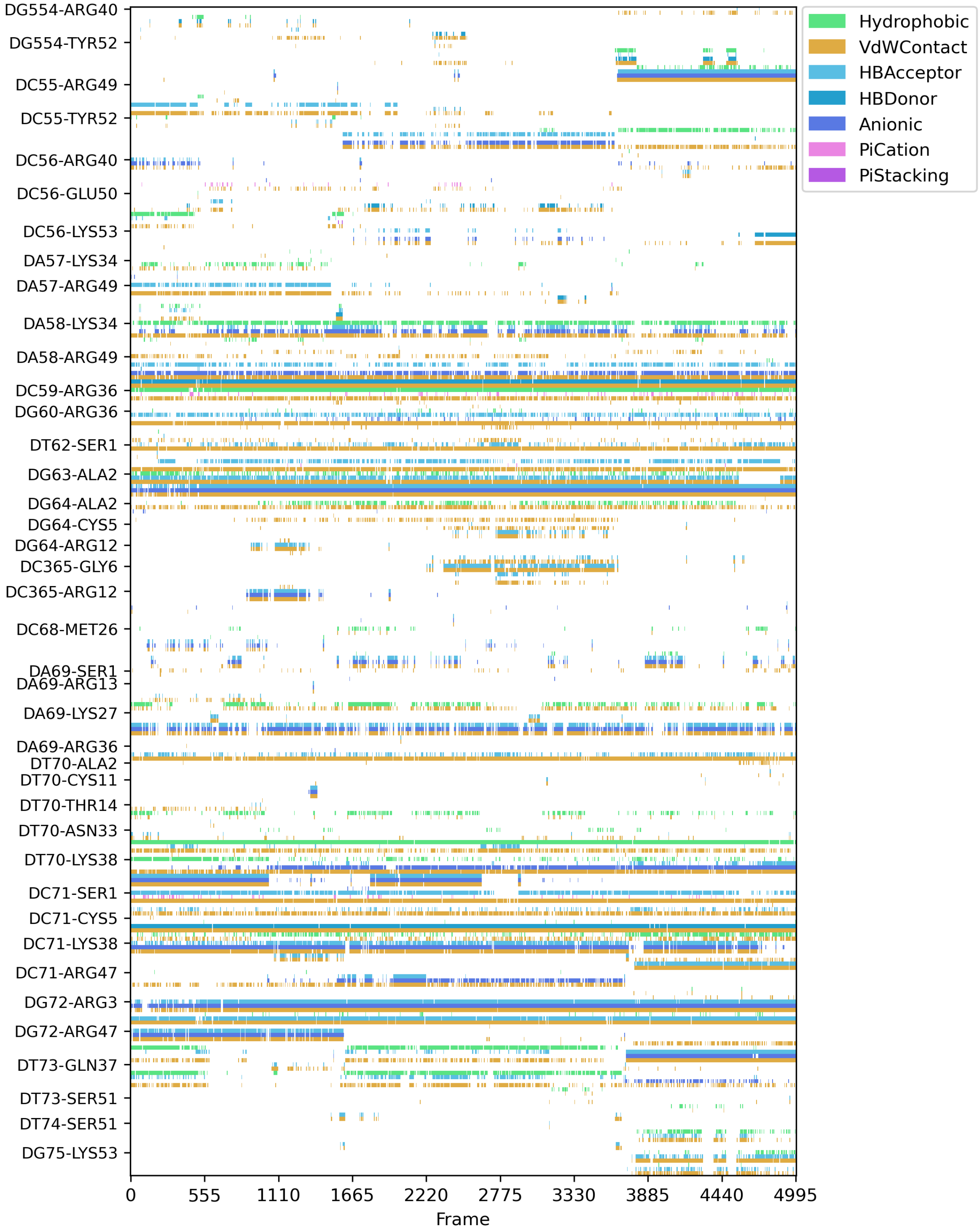


Figure S12: Bar plot depicting the interaction fingerprint between the protein and DNA in CFP1-CpG Complex (Replica 4).

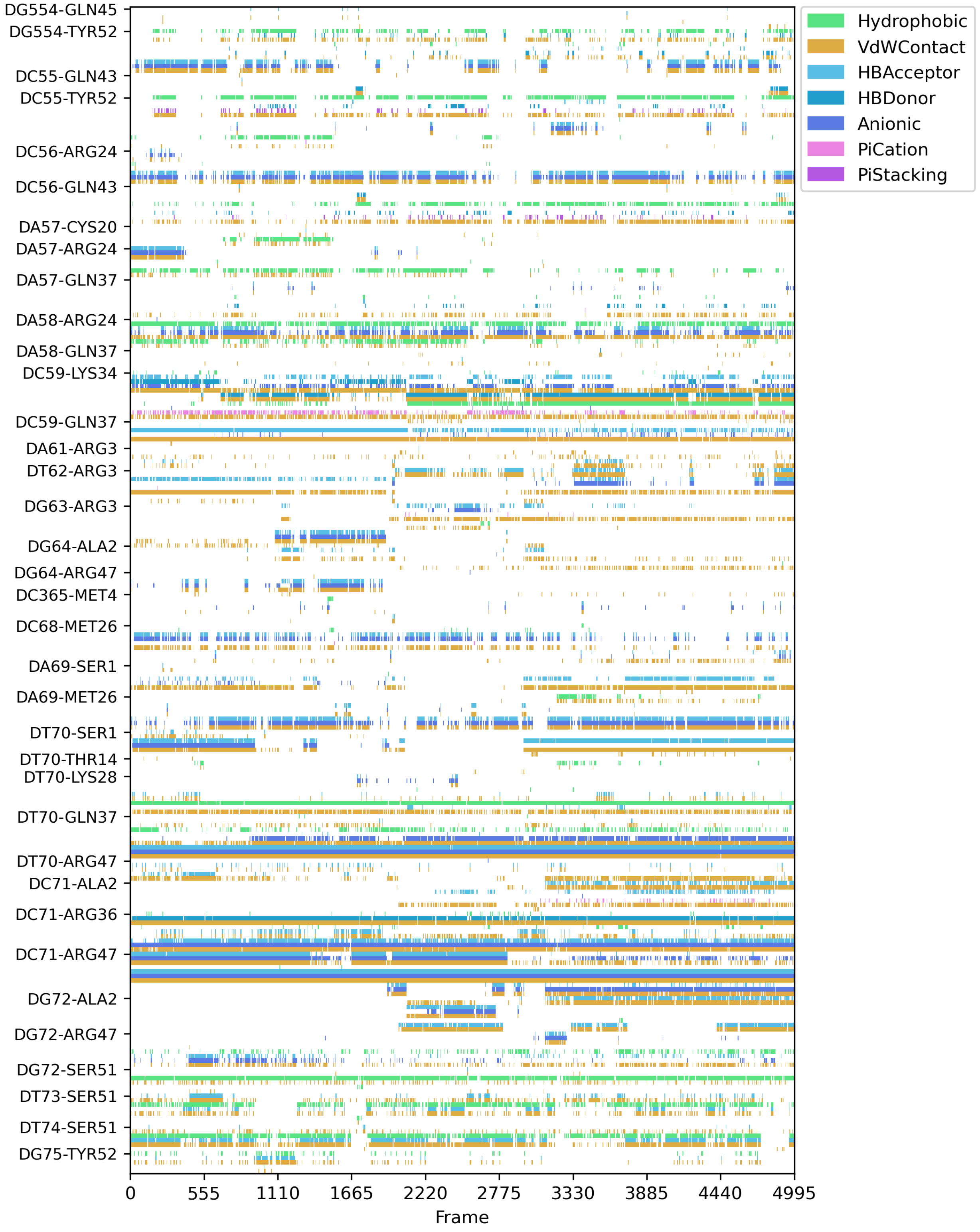


Figure S13: Bar plot depicting the interaction fingerprint between the protein and DNA in CFP1-CpG Complex (Replica 5).

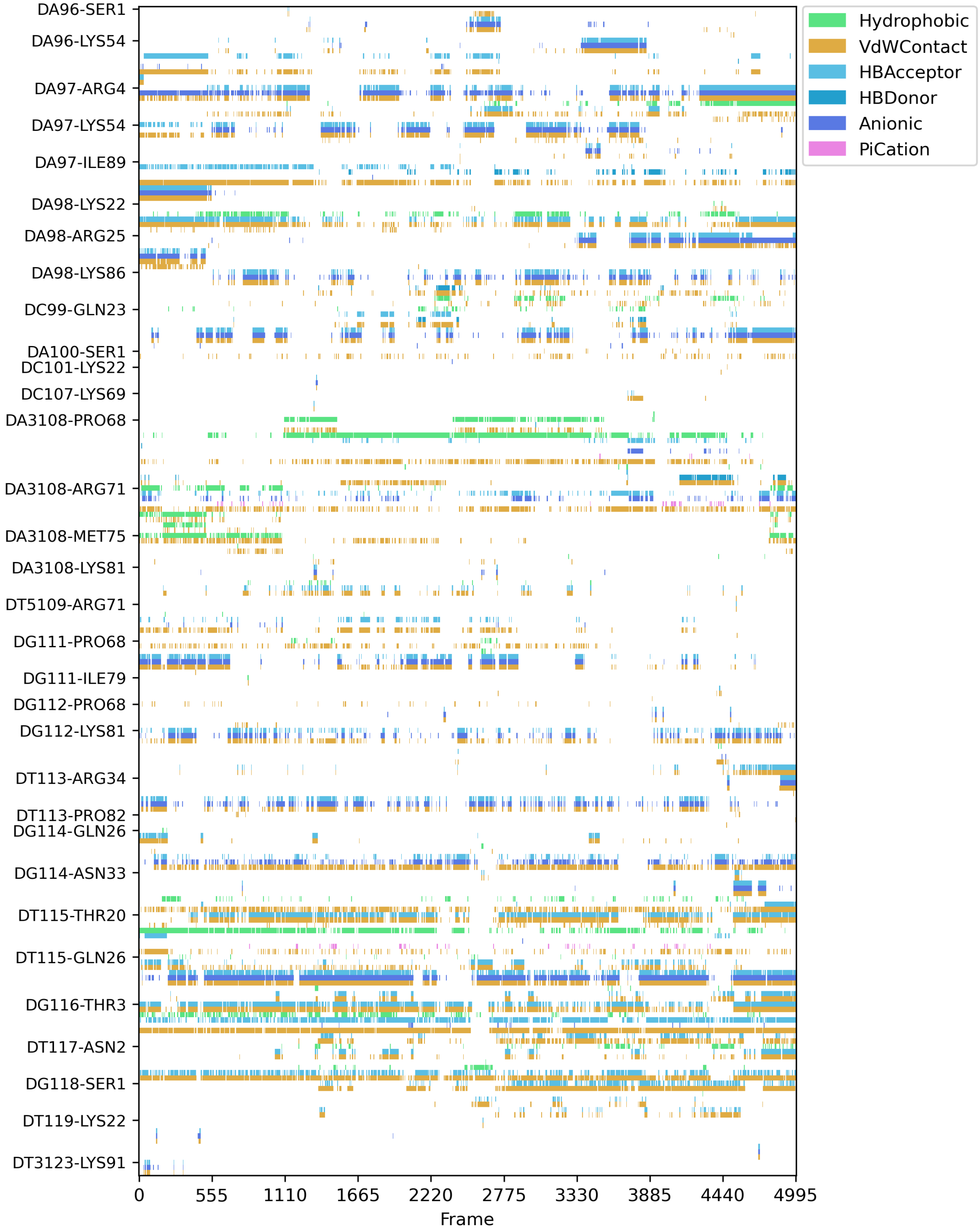


Figure S14: Bar plot depicting the interaction fingerprint between the protein and DNA in MC1-DNA Complex (Replica 1).

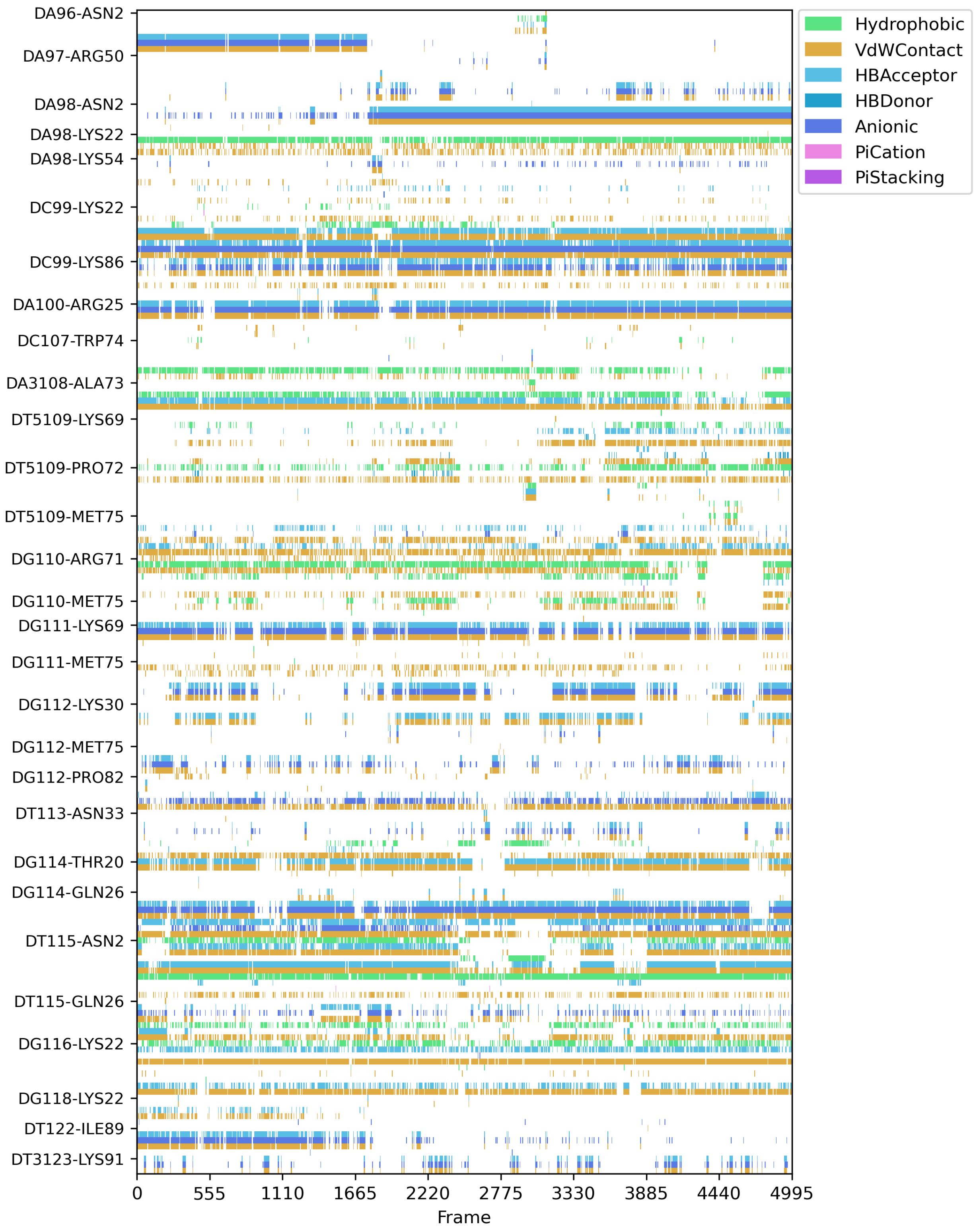


Figure S15: Bar plot depicting the interaction fingerprint between the protein and DNA in MC1-DNA Complex (Replica 2).

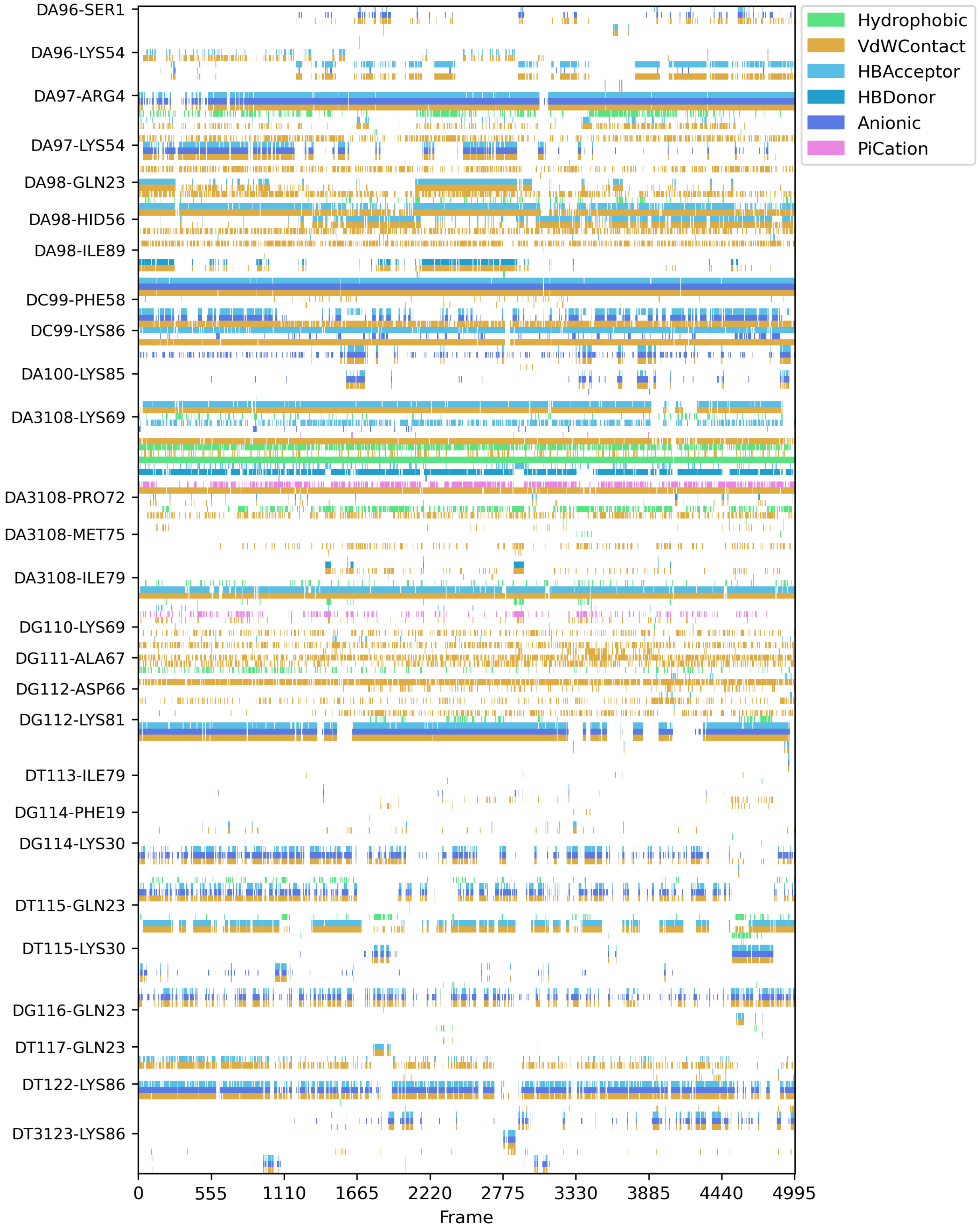


Figure S16: Bar plot depicting the interaction fingerprint between the protein and DNA in MC1-DNA Complex (Replica 3).

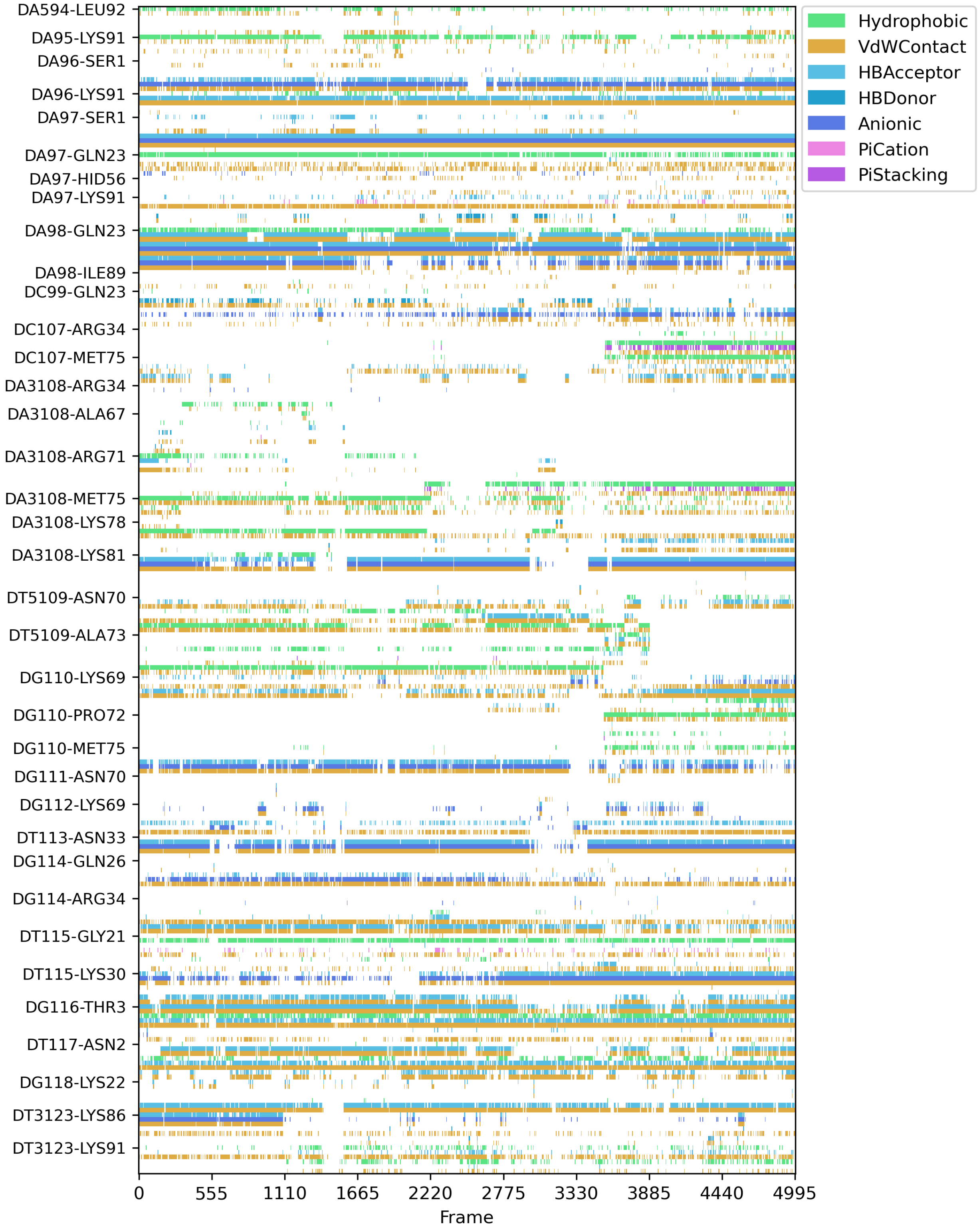


Figure S17: Bar plot depicting the interaction fingerprint between the protein and DNA in MC1-DNA Complex (Replica 4).

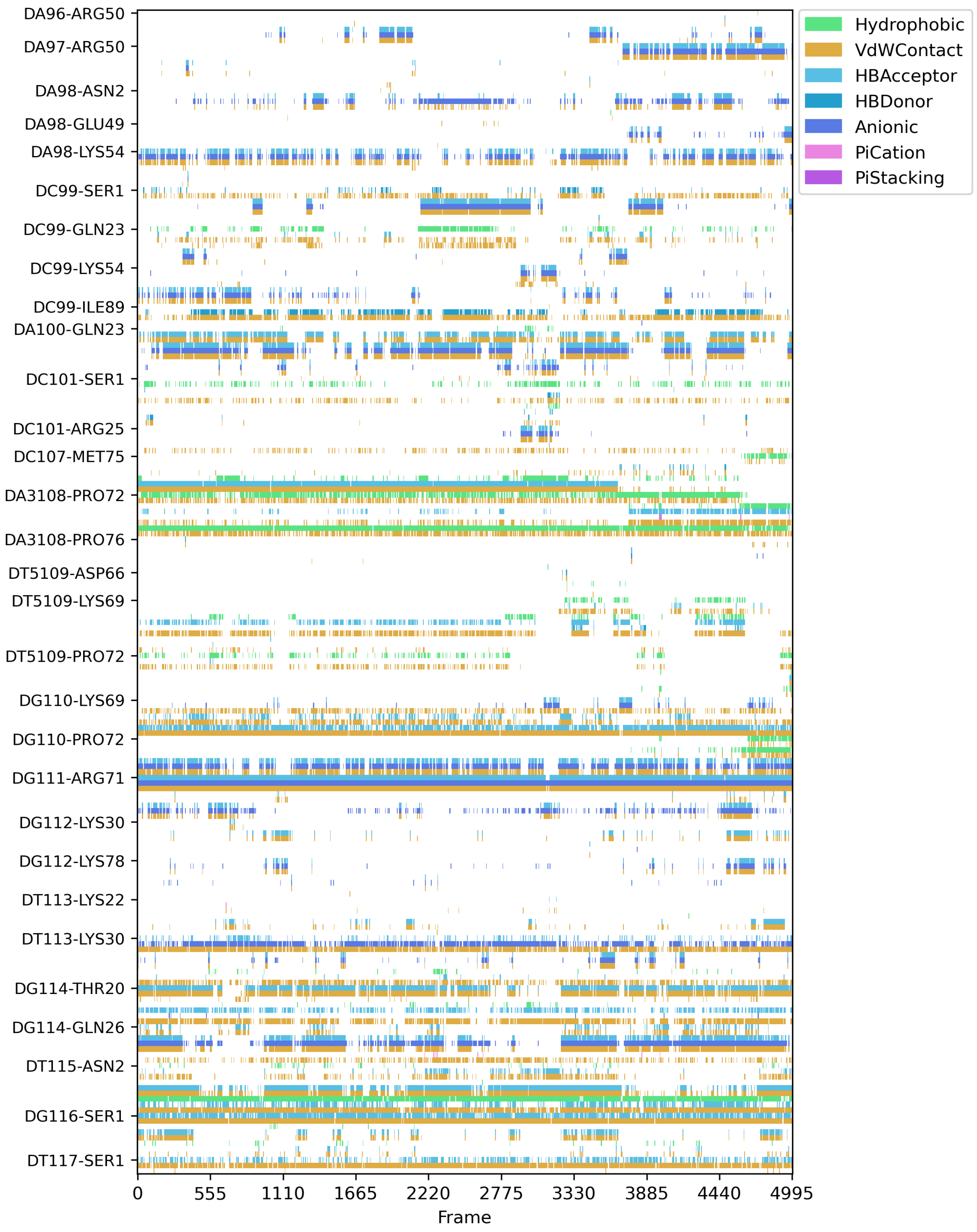


Figure S18: Bar plot depicting the interaction fingerprint between the protein and DNA in MC1-DNA Complex (Replica 5).

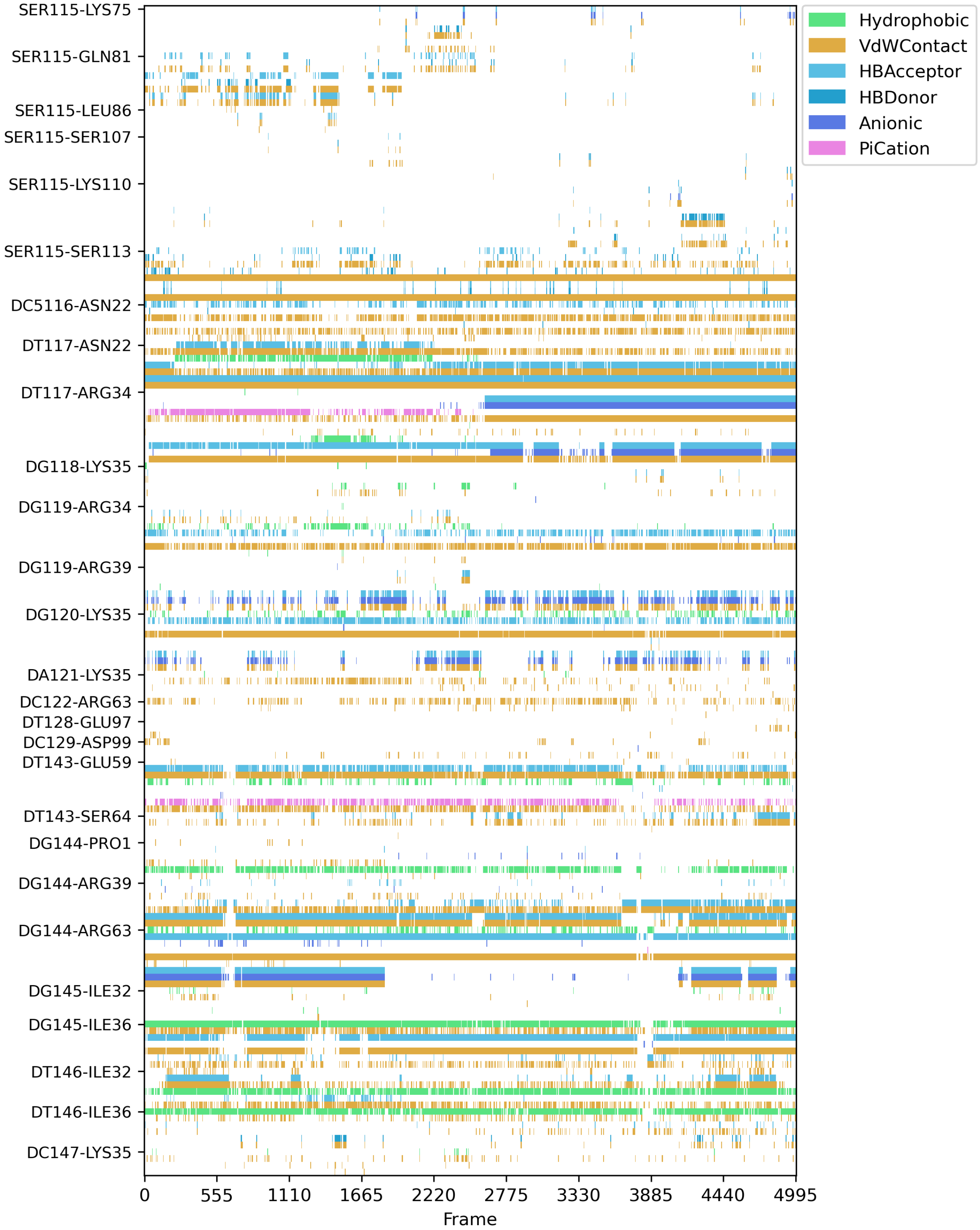


Figure S19: Bar plot depicting the interaction fingerprint between the protein and DNA in SopB-DNA Complex (Replica 1).

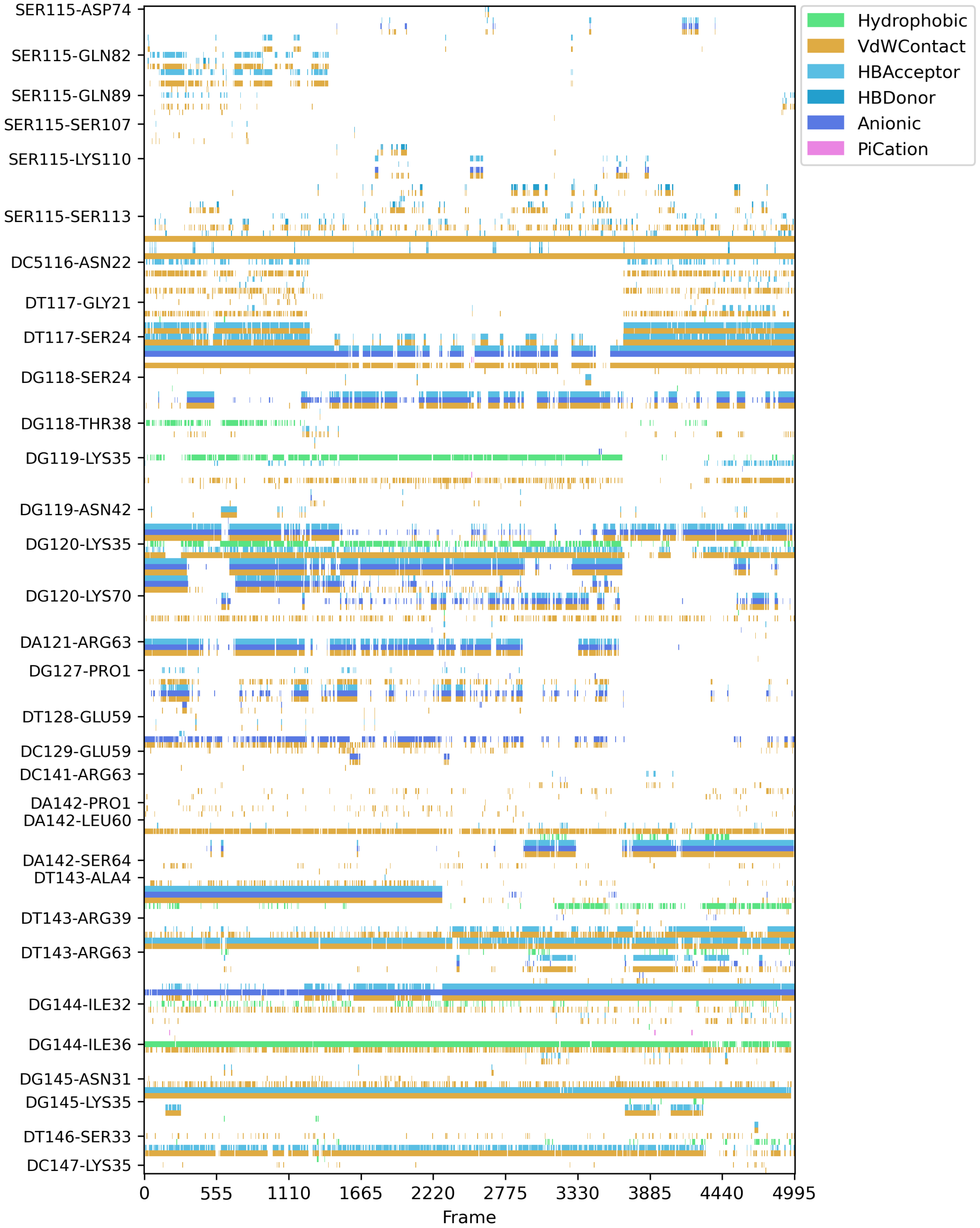


Figure S20: Bar plot depicting the interaction fingerprint between the protein and DNA in SopB-DNA Complex (Replica 2).

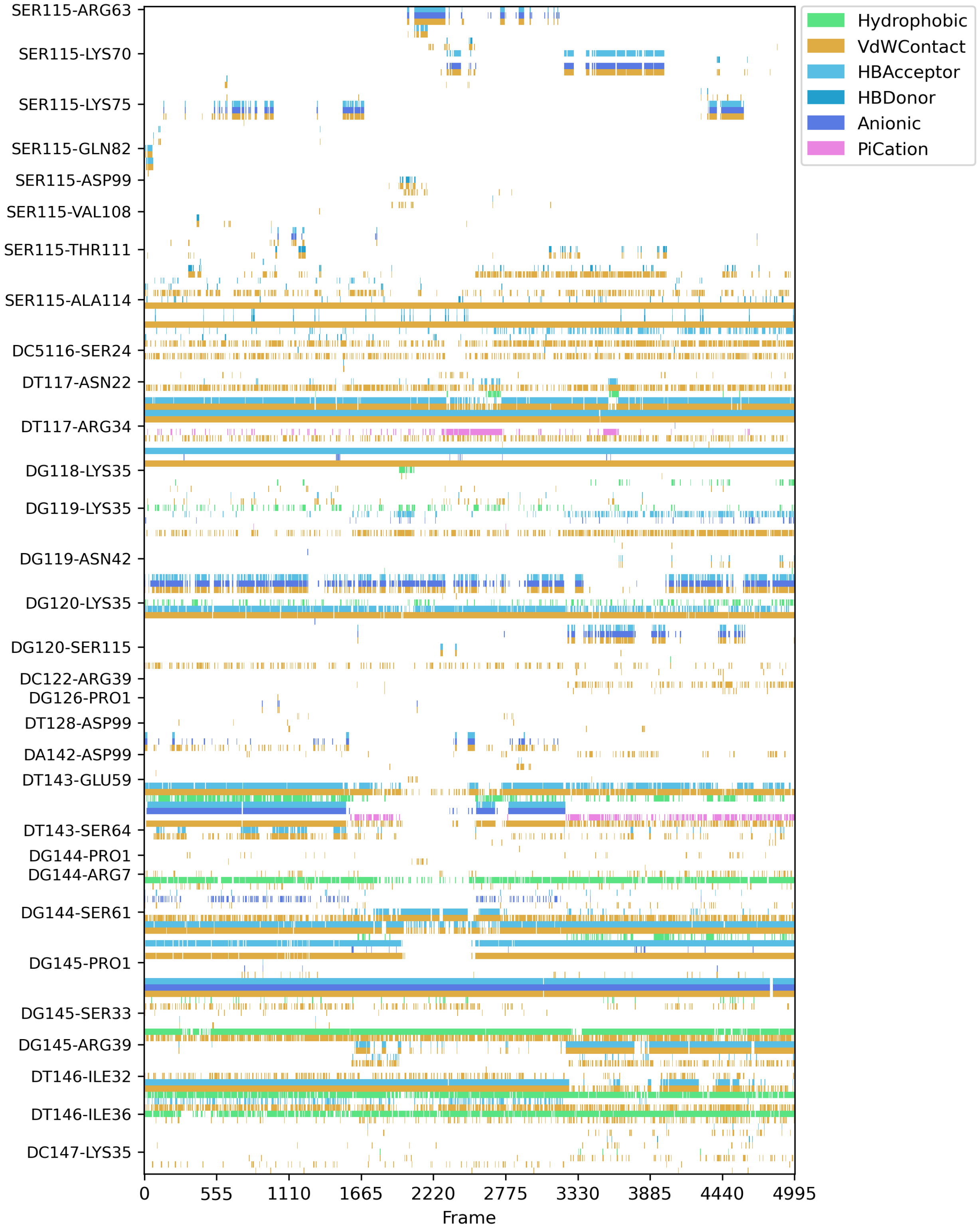


Figure S21: Bar plot depicting the interaction fingerprint between the protein and DNA in SopB-DNA Complex (Replica 3).

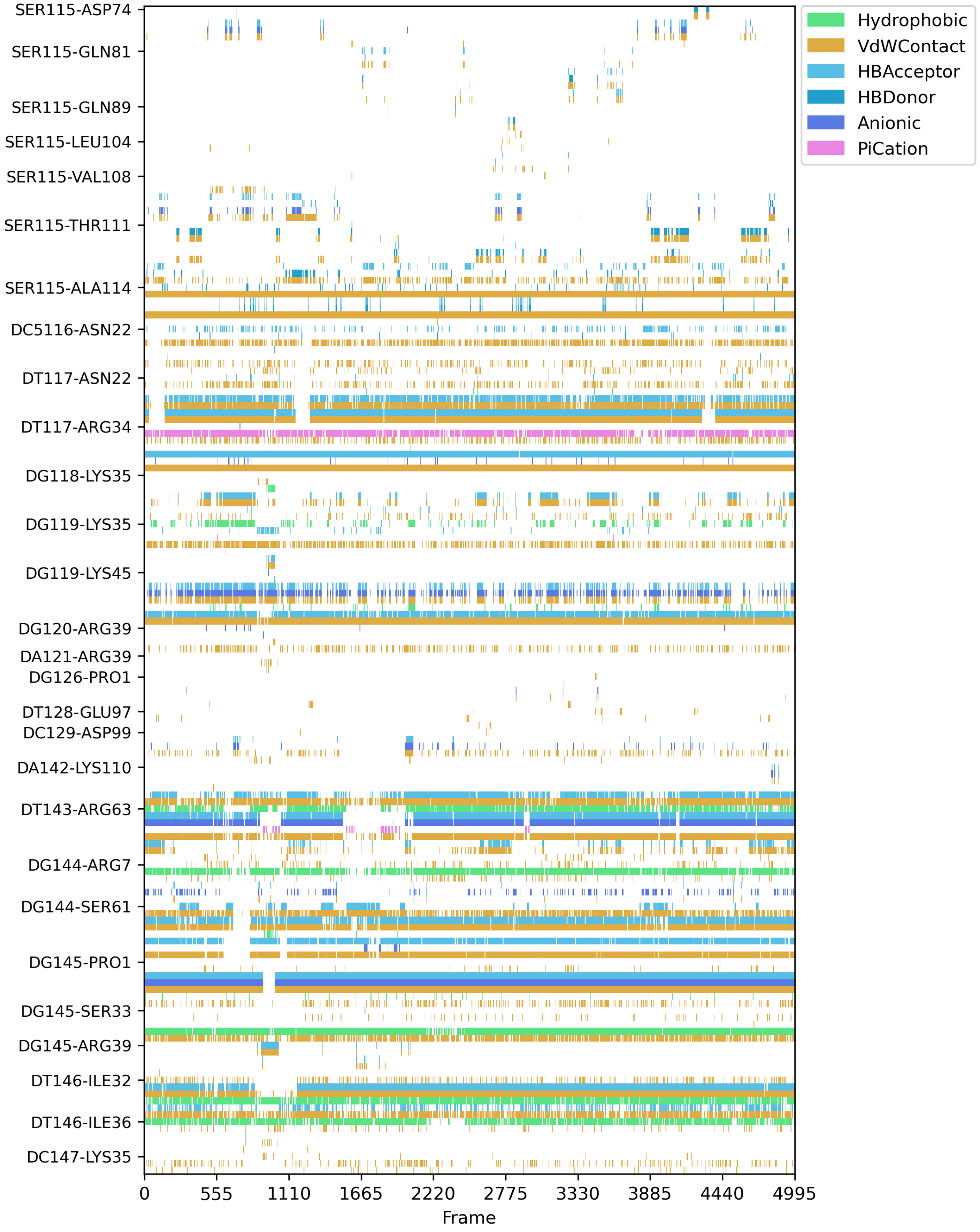


Figure S22: Bar plot depicting the interaction fingerprint between the protein and DNA in SopB-DNA Complex (Replica 4).

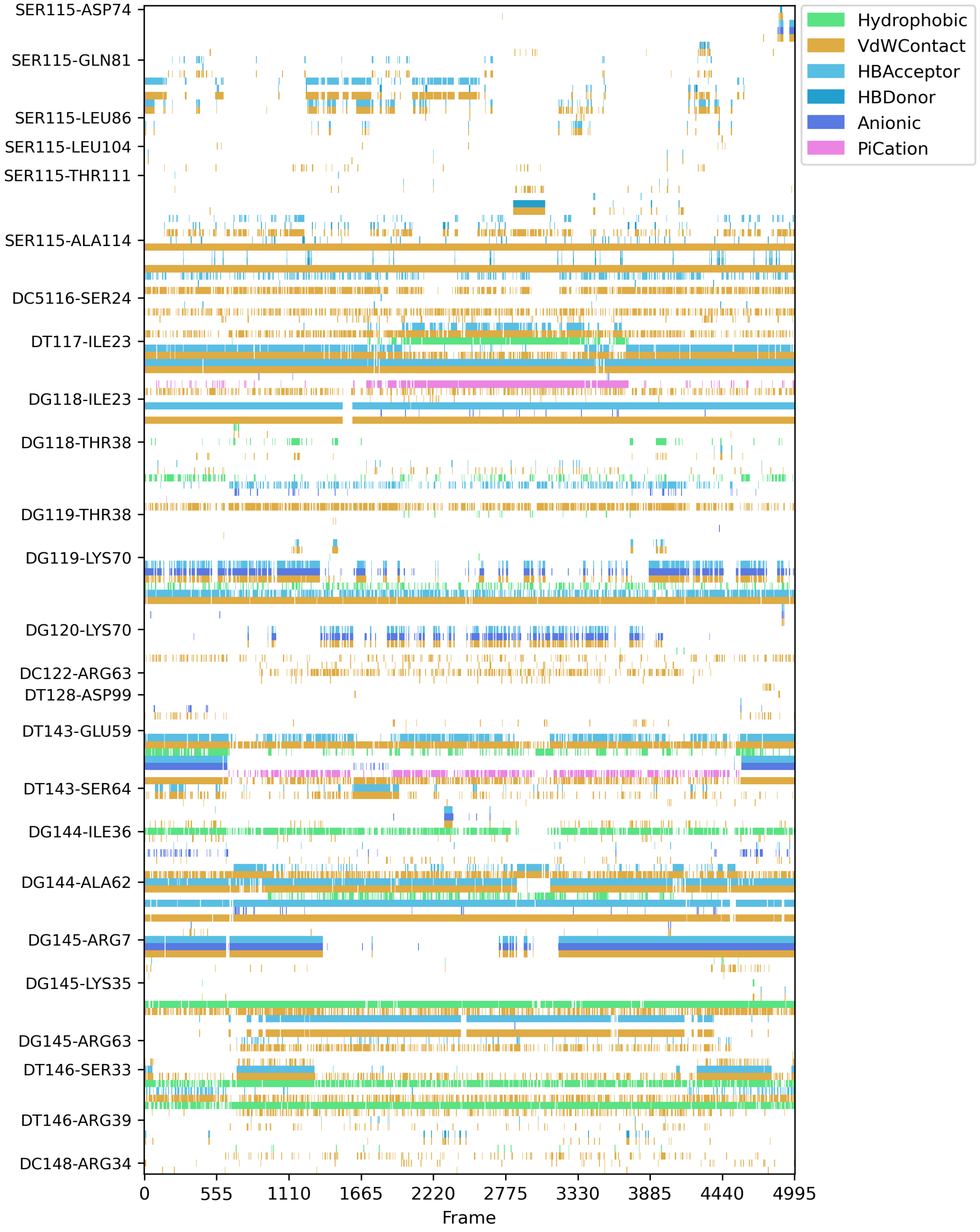


Figure S23: Bar plot depicting the interaction fingerprint between the protein and DNA in SopB-DNA Complex (Replica 5).
